# Mechanisms and control of a novel vocalization: The singing mouse song is a whistle that depends on air sac inflation

**DOI:** 10.1101/2025.05.16.654575

**Authors:** Samantha K. Smith, Jonas Håkansson, Paul W. Frazel, Michael A. Long, Coen P.H. Elemans, Steven M. Phelps

## Abstract

Identifying variation in vocal morphology and sound production mechanisms is essential to understanding vocal diversity. Rodents provide an ideal system for exploring this variation as they use multiple sound production mechanisms and have novel vocal structures whose morphology varies interspecifically. Here, we describe the laryngeal morphology and identify the sound production mechanism of Alston’s singing mouse (*Scotinomys teguina*), which produce stereotyped songs. We used micro-computed tomography to examine laryngeal morphology and manipulated excised larynges and surgically ablated a laryngeal muscle to determine sound production mechanism and frequency control. Laryngeal manipulations indicated that a whistle mechanism, likely an edge tone or shallow cavity, produces song. Singing mouse whistles are unique compared to other rodents because they rely on the inflation of an enlarged intralaryngeal air sac called the ventral pouch. Whistle frequency can be controlled by ventral pouch inflation, laryngeal airflow, and by cricothyroid muscle action. Cricothyroid ablation inhibited frequency modulation *in vivo*, suggesting that singing mice use this muscle during singing. Singing mouse laryngeal morphology and vocal mechanism are distinct from other Neotomids; instead of using vocal fold oscillations for loud, long-distance calls and whistles for close-range interactions, singing mice appear to use whistles for distant and close exchanges by inflating their intralaryngeal air sac. Air sacs have evolved repeatedly among vocalizing mammals and our results indicate a new role for these structures in generating sound. Together, our results expand on an emerging story of how biomechanic and morphological variation contributes to vocal diversity.

## Introduction

Understanding how complex structures take on new functions is a central goal of biology^1,2^. In the tetrapod larynx, for example, structural innovations that enable vocalization are constrained by the demands of respiration and airway protection^3^, offering a case study in how complex structures acquire new functions while maintaining others. Despite these constraints, mammals use their larynx to make a tremendous diversity of vocalizations, including the infrasonic calls of elephants^4^, the ultrasonic squeaks of mice^5^, the frequency-modulated pulses of long-legged bats^6^, the grunts of male reindeer^7^, and the long-distance chirps of cheetahs^8^. To address how the constrained larynx gives rise to such a broad range of acoustic outputs, we investigated the laryngeal mechanisms underlying a unique vocal innovation, the advertisement song of Alston’s singing mouse (*Scotinomys teguina*).

Rodents represent a particularly promising group for exploring vocal variation because they exhibit novel laryngeal morphology^9^ and use multiple sound production mechanisms^10–14^. Most tetrapod vocalizations are generated by airflow-induced tissue vibration, known as the myoelastic-aerodynamic theory of phonation (MEAD)^15,16^. These include laryngeal vocalizations of anurans and mammals, as well as the syringeal vocalizations of birds^16^. However, many rodents produce high-frequency sounds using a distinct mechanism involving whistle-like air vibration^10,11,14,17^. Recent work in a few rodent genera has revealed important links between laryngeal anatomy, vocal mechanism, and vocal output^14,18,19^. Expanding this work to species with divergent vocalizations like singing mice, offers opportunities to discover the anatomical and physiological innovations that support vocal complexity.

We use Alston’s singing mouse, *Scotinomys teguina*, to examine how differences in laryngeal mechanisms and morphology contribute to acoustic diversity. Singing mice are diurnal rodents that live in the montane grasslands of central America^20,21^. They make long, highly stereotyped songs consisting of rapidly-repeating, tonal, frequency-modulated notes that span ∼30 kHz in as little as 12 msec^22–27^ and reach sound pressure levels of ∼60 dB at 1 meter (Figure 1). Singing mice use songs for mate-attraction and in male-male competition. In these interactions, songs are rapidly exchanged between individuals (counter-singing) at speeds resembling human conversational speech^28^. Songs also exhibit individual- and population-level variation^27^. Like other vocalizing species, song spectral features are repeatable within individuals, and are thought to be limited by morphology; these song features are more heritable than song-effort components in wild-caught singing mice^29^.

**Figure 1.**
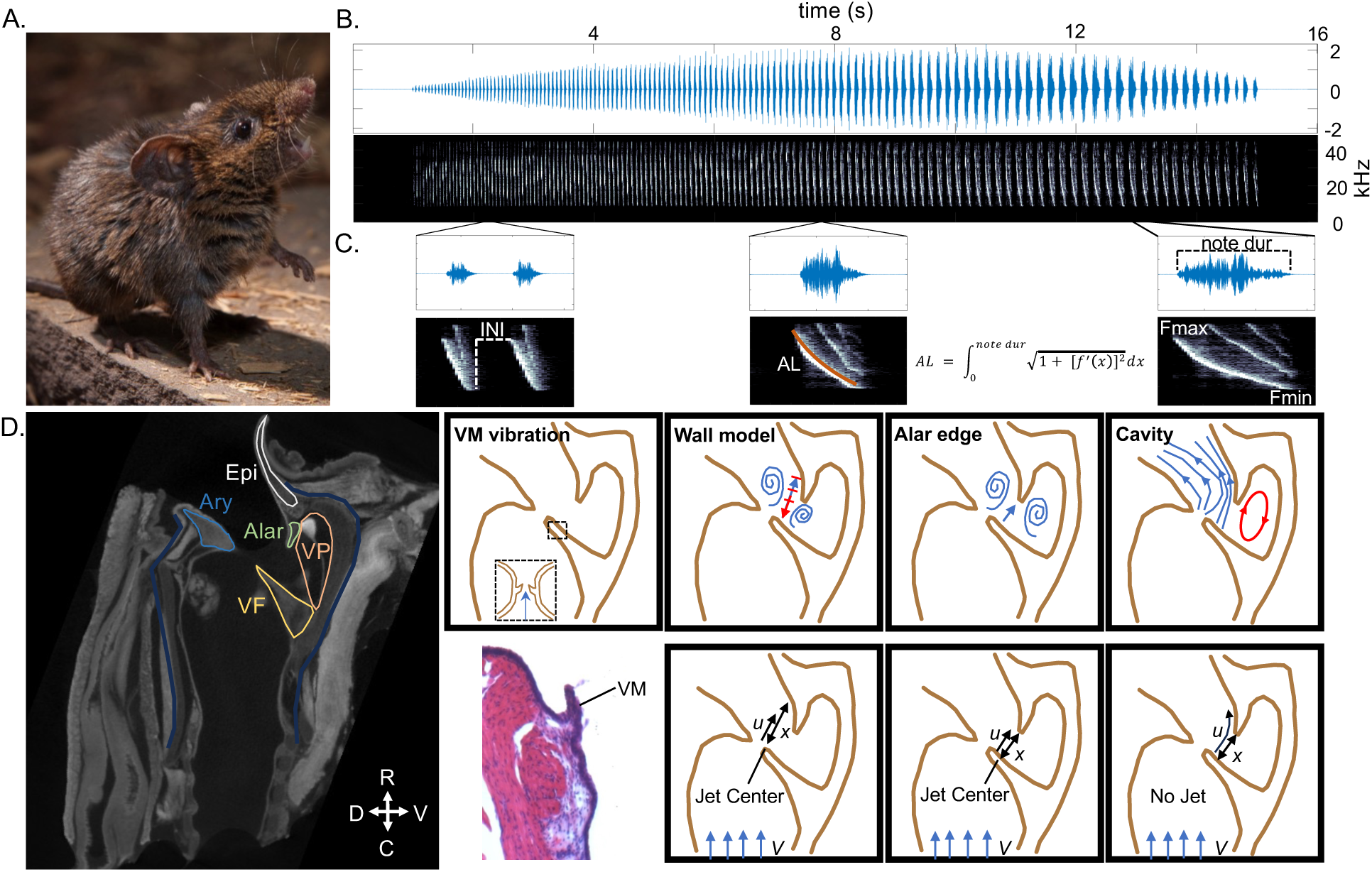
*Scotinomys teguina* song and potential sound production mechanisms. (A) Male *S. teguina* singing. (CC) Christopher Auger-Dominguez. (B) Oscillogram (top) and spectrogram (bottom) of a typical male song. (C) Oscillograms and spectrograms of 0.15 sec of song from the beginning (left), middle (center), and end (right). (D) (left) Mid-sagittal view of an iodine-contrasted, μCT-scanned larynx. (right bottom) H&E-stained, transverse section of vocal folds with vocal membrane (VM). (right) Four potential song production mechanisms. Top plots show air flow (blue arrows) and feedback (red arrows). Bottom plots show variables that control frequency. u = air speed, x = distance air travels, V = air velocity. Ary = arytenoid cartilage, Epi = epiglottis cartilage, Alar = alar cartilage, VP = ventral pouch, VF = vocal folds. R = rostral, C = caudal, D = dorsal, V = ventral. INI = internote interval, AL = arc length, note dur = note duration, Fmax = frequency maximum, Fmin = frequency minimum.

Singing mice are a part of the Neotominae subfamily noted for their audible and loud vocalizations, a rare trait among small rodents. Singing mice and their close relatives (Baiomyini) produce long, frequency-modulated vocalizations with higher maximum frequencies, greater average bandwidths, and more rapid pulse rates than other Neotomids^22,30,31^. Among Baiomyini, the Alston’s singing mouse song is the most extreme (e.g., *S. teguina*: ∼100 notes/song, 10 s duration; *B. taylori*: ∼20 notes/song, 2 s duration). The unique nature of the singing mouse song compared to other rodents provides opportunities to identify how laryngeal morphology and sound production mechanisms are modified to generate complex vocalizations.

Although the production of song and its social exchange appear to be precisely controlled in singing mice, we know little about the mechanisms that generate sound and control frequency. Note frequencies span 43 to 10 kHz (Figure 1) which could be consistent with vocal membrane vibration or whistle mechanisms. Histological characterization of singing mouse larynges indicates that they have the morphological features required for vocal membrane and whistle mechanisms, including an enlarged air sac called the ventral pouch^32^. Interestingly, the louder, long-distance calls of other Neotomids are produced with tissue vibration, while close-range USVs are whistles^12,18,19^. However, *Baiomys* use a whistle mechanism to produce both long- and close-range calls^13^.

Here, we examined whether singing mice use vocal membrane vibration or a whistle mechanism to sing. We first used diffusible, iodine-based, contrast-enhanced, micro-computed tomography^33–35^ to describe the 3-D anatomy of the singing mouse larynx. To identify song production mechanism and examine frequency control, we studied the excised larynx while recording audio and high-speed video. We then determined the effect of ablating the cricothyroid (CT) on singing *in vivo*. We chose this muscle based on its activity during vocal fold and whistle sounds in other rodents, as well as its general role in frequency modulation across tetrapods. We found that the singing mouse song is an aerodynamic whistle that requires inflation of an intralaryngeal air sac called the ventral pouch. Air sacs have evolved many times in vocalizing mammals and our results highlight a novel role for these structures in sound generation.

## Methods

All animal procedures were approved by UT Austin and NYU Medical IACUC. µComputed Tomography (µCT) *Scotinomys teguina* (n=6, 3 males, 3 females) were taken from the Phelps laboratory colony at UT Austin. The colony was outbred from a wild population caught at Quetzal Education Research Center (QERC) in San Gerardo de Dota, Costa Rica. All mice were euthanized with an overdose of isoflurane. We removed the larynges, drop fixed them in 4% PFA at 4°C for 24 hours, dehydrated them with a series of ethanol incubations (70%, 95%, 100%) over 36 hours, and then stained them with 2.5% Lugol’s iodine for 16 hours.

University of Texas High-Resolution X-ray CT facility scanned larynx samples using an Xradia MicroXCT 400 system with a 4X objective (80kV, 10W). Detector and source distances were optimized for all samples (detector 8 mm, source -47 mm). The end reference was 45 frames at 2s each. Images were reconstructed (Xradia Reconstructor, Table S1). Each sample had a voxel size of 5.75 µm. See Table S2 for XYZ values and total number of slices for each larynx sample.

After image acquisition, we reconstructed the data using ImageJ (v. 2.3.0/1.53q) and Avizo (v. 2022.2). We segmented the data to identify all cartilages, 5 muscle bodies (cricothyroid, lateral thyroarytenoid, medial thyroarytenoid, posterior cricoarytenoid, lateral cricoarytenoid), the interior airway, and the ventral pouch. After segmentation, we visualized volumes using Avizo and ImageJ.

### Excised larynx manipulations

15 singing mice (10 males, 5 females) were euthanized with an overdose of isoflurane. Larynges were extracted and frozen on dry ice and stored at -80C. We sourced these animals from the Phelps (UT) and Long (NYU) laboratories. Prior to euthanasia, mice were recorded for spontaneous vocalizations at the Long laboratories (NYU) in recording chambers (Med Associates) using condenser microphones (Avisoft Bioacoustics CM16/CMPA) sampled at 250 kHz (digitized with Avisoft UltraSoundGate 116Hb)^28^.

We thawed each larynx at 4°C, and micro dissected it in Ringer’s solution to remove all extra-laryngeal tissue. The larynx was then mounted on the experimental rig^36^ in the Elemans Lab (SDU), which consisted of an airflow hose connected to a mounting base where the larynx sits, micromanipulators to adduct arytenoids and glottis, a high-speed camera (MotionPro X4-M-4, Integrated Design tools, Inc., USA, 250 fps) placed above the mounting base, a ¼-inch pressure microphone-preamplifier assembly (model 46BD, frequency response ± 1 dB 10 Hz-25 kHz and ± 2 dB 4 Hz-70 kHz G.R.A.S., Denmark), and a servo motor (Aurora Scientific model 300c). Subglottal air pressure and servo motor action could be controlled with a custom Matlab graphical user interface, while simultaneously recording audio and video of larynges.

In the first experiment, we assessed whether tissue or air vibration produced frequencies within the range of the singing mouse song and identified the structures necessary for sound production *in vitro*. We removed the superior portion of the epiglottis, so that we could visualize the glottal opening and access the arytenoid cartilages. We first forced air through the larynx (pressure ramp 0-5 kPa, speed 1 kPa/sec) without adducting the larynx, recording video and audio. Then we used the micromanipulators to press on the arytenoids and close the glottis, approximating a phonation position and repeated the airflow protocol, recording sound and taking video. After this, we repeated the airflow protocol with the following manipulations: (1) filling the ventral pouch with wax or metal balls and (2) removing the alar cartilage.

We then assessed how imitating cricothyroid action and changing airflow impacted frequency *in vitro.* We placed a suture on the superior edge of the medial thyroid. We attached the suture to a servo arm and placed a small, mirrored prism next to the larynx to capture the top and side view. We ran a custom step function where air passed through the larynx at an initial pressure of 5 kPa and was reduced to 0.5 kPa by 0.5 kPa/second. The servo pulled on the thyroid cartilage once per second, imitating the action of the cricothyroid muscle^37,3^ at each air pressure. We then cut away the alar cartilage and reran the custom step function.

Data were analyzed using Matlab R2020a (9.8.0.1323502). We used a high-pass Butterworth filter on sound data, calculated spectrograms (nfft=1024, overlap=0%, flat top window), used ridge detection to identify the dominant frequency, and then used Weiner’s entropy^38,39^ to differentiate between signal and noise (threshold 0.4). Weiner’s entropy tends to do a good job detecting the fundamental frequency in our dataset.

However, entropy is inflated when there is a lot of frequency modulation. Where frequency rapidly changed, we extracted fundamental frequency using the Yin algorithm^40^. Yin is powerful because it detects fundamental frequency but becomes unreliable with frequencies that surpass ¼ of the sampling rate^40^. We kept Yin-detected frequencies if power > 0.085 and aperiodicity < 1. Using both methods allowed us to accurately extract all sounds in the dataset. We then measured relevant parameters including dominant frequency, sound pressure level, subglottal air pressure, tracheal airflow, and sound amplitude.

Videos were rotated such that all movement occurred along the x- and y-axis. We selected 3-4 videos and used kmeans clustering to extract 15 frames per video with DeepLabCut^41,42^ (v. 2.3). For all videos, we applied landmarks along the left and right edges of the glottal opening, the alar and epiglottis cartilages (Table S3). We placed additional landmarks at the thyroid edge to videos from the second experiment. We used these frames as training data for a deep learning algorithm. This procedure was done for each larynx separately. See the supplemental text for details on the deep learning algorithm we used (Figure S1, Table S4). We applied a 4^th^ order, zero-lag, low-pass, Butterworth filter (cutoff frequency = 10) on resulting landmark coordinates, defining the center of the FFT window as the length of time between video frames. We used these coordinates to track the movement of the ventral pouch and alar cartilage. Since ventral pouch inflation occurs along the z-axis with respect to the camera, we used the movement of the alar edge and epiglottal stems as a proxy for inflation. We then aligned the movement data with the sound data to identify the position of the larynx at sound initiation and determine the relationship between tissue movement and frequency.

### Cricothyroid muscle ablation and recording

10 male singing mice underwent surgery (5 sham, 5 ablation) and audio recording to identify the role of the cricothyroid muscle in vocalization. Prior to surgery, each animal was recorded for spontaneous vocalizations over a 24-hour period using ACO Pacific microphones on RX8 hardware sampled at 97.6 kHz. One week after recording, we anesthetized each male using isoflurane gas and made an incision on the ventral neck. We cut away a small portion of the sternohyoid and sternothyroid to reveal the cricothyroid. The location of the cricothyroid was confirmed by identifying the thyroid and cricoid cartilages which are superior and inferior to this muscle respectively^37,3^. Five animals received sham surgeries, where the surgeon closed the incision after revealing the cricothyroid. The other 5 animals received ablation surgeries, where the surgeon bilaterally cut away as much of the cricothyroid as could be accessed. Animals recovered for 3 days before being recorded for spontaneous vocalizations over a 24-hour period.

At the end of the 24-hour period, we euthanized the animals using CO_2_ and immediately dissected out the larynx. We fixed the larynges at 4°C overnight in 4% paraformaldehyde in 1X phosphate buffered saline (pH 7.4), cryoprotected them with sucrose (15% overnight, 30% for 7 hours), and then froze larynges in OCT compound. We took every other 20-µm thick section on a NX50-2 cryostat at the Center for Biomedical Research Support Microscopy and Imaging Facility at UT Austin (RRID:SCR_021756). All slides were stained with Hematoxylin and Eosin (H&E kit, Abcam, ab245880). We imaged slides on a Leica DMIL microscope with a ProgRes CFscan camera (1360 x 1024 pixel resolution) and CapturePro 2.7.6 software at 5x objective. We merged overlapping images into a single image of each section using photoshop.

We estimated the volume of intact cricothyroid muscle using ImageJ (v. 2.3.0/1.53q). Lateral cricoarytenoid and thyroarytenoid muscles were also measured because we found that the ablation impacted other muscles in some samples. To estimate intact muscle volume, we measured the area of non-ablated muscle in all sections using the polygon tool (imageJ v. 2.3.0/1.53q). We then calculated the volume using the equation^43^:

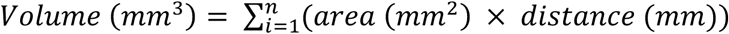

### Song Analysis

We used matlab scripts (https://github.com/samksmith/steg_muscle_ablation) that automatically extracted songs from the audio recording and took whole song and individual note measures. Each note of the singing mouse song consists of a downward frequency sweep that can be described by a quadratic equation^27,44^. We calculated the arc length of each note using the equation:

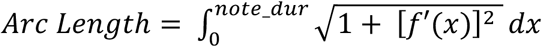

where *f*(*x*) = *FMa* ∗ *x*^2^ + *FMb* ∗ *x* + *FMc*

We generated and visually-inspected spectrograms and oscillograms for all songs using Matlab (R2020a, v. 9.8.0.1323502). All quantitative analyses were done in R (v. 4.1.2). Because we expected that singing mice who received an ablation would have a reduced capacity to modulate frequency, we examined mean note-by-note arc length values. Song length varied between and within individuals (although surgical groups did not differ in the number of songs they sang across the two timepoints, see Figure S2). Because arc length changes consistently across song, we divided each arc length by its note duration and compared normalized arc length across timepoints for each individual. In addition to individual level analyses, we fit a series of linear mixed-effects models (4 random intercepts, 1 random slope model) to identify whether treatment and timepoint explain variation in arc length (lme4^45^, v. 1.1-27.1). We did likelihood ratio tests (lmtest::lrtest^46^, v. 0.9-40) and used AIC model selection (AICcmodavg::aictab^47^, v. 2.3-1) to identify the best model. Finally, we used the MuMIn package (v. 1.46.0) to calculate the marginal and conditional R^2^ for the best model. We ran the same models on a reduced dataset where individual A was excluded. This individual received an ablation surgery which damaged other intrinsic muscles. Finally, we identified whether estimated muscle volume predicts the change in mean, normalized arc lengths using linear modeling (stats::lm, v. 4.1.2). We created 7 models (6 combinations of muscles volumes and 1 null model) and used a likelihood ratio test and AIC model selection approach to identify the best model.

## Results

Here, we examined whether singing mice use vocal membrane vibration or an aerodynamic whistle mechanism to produce song.

### 3-D laryngeal anatomy

We first used diffusible, iodine-based, contrast-enhanced, micro-computed tomography^33–35^ to describe the 3-D anatomy of the singing mouse larynx. As previously reported^48^, singing mice have a ventral pouch as well as the same cartilages (cricoid, thyroid, arytenoids, alar, epiglottis) and intrinsic musculature (cricothyroid, thyroarytenoid, lateral cricoarytenoid, posterior cricoarytenoid, transverse and oblique arytenoids) as other rodents. We also found that the alar and epiglottis cartilages are fused in singing mice, as reported in northern pygmy mice (*Baiomys taylori*) ^13^ (Figure 2).

**Figure 2.**
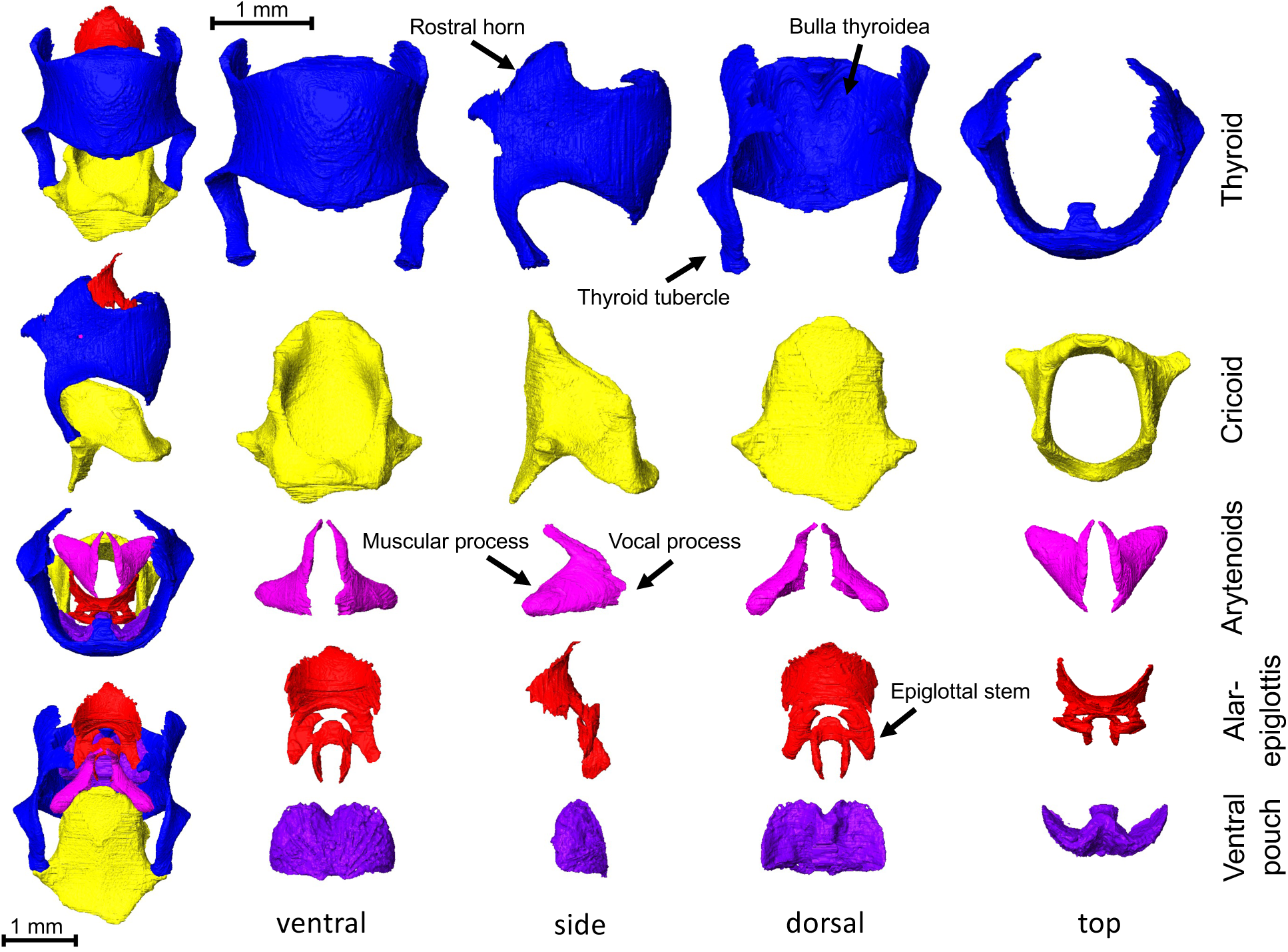
Singing mouse larynx reconstructions. From top to bottom, four views of the thyroid (blue), cricoid (yellow), arytenoids (pink), alar-epiglottis (red), and ventral pouch (purple). (Left column) From top to bottom, the ventral, side, top, and dorsal views of the 5 laryngeal structures together. The thyroid, cricoid, arytenoids, alar, and epiglottis are cartilages. The ventral pouch is an air sac positioned between the thyroid and alar cartilages. The alar and epiglottis are fused at two points along the alar edge on the ventral side. The ventral pouch extends medio-laterally and dorso-ventrally.

#### Cricoid cartilage

The cricoid cartilage forms the lower portion of the larynx and is connected to the thyroid cartilage. The dorsal plate is broad while the ventral plate is narrow. Its ring-like structure forms a rectangular opening that appears longer along the dorso-ventral axis than that of the medio-lateral. There is a protuberance on the narrow ventral surface where the cricothyroid muscles insert.

#### Thyroid cartilage

The thyroid cartilage is positioned superior to the cricoid where it connects via prominent thyroid tubercles. The ventral face of the thyroid is elongate along the rostro-caudal axis. The rostro-ventral margin bends dorsally in singing mice. On the inner wall of the thyroid, an indent forms the bulla thyroidea, where the ventral pouch is located. The caudal edge of the ventral pouch is formed by a small protuberance, where the vocal folds attach to the inner wall of the thyroid. The rostral horns are broad, but angle steeply downward in singing mice.

#### Arytenoid cartilages

The arytenoids sit dorsal to the thyroid and attach to the vocal folds ventrally. The vocal process, which makes up part of the glottis, is short, while the muscular process is long.

#### Alar-epiglottis cartilage

The epiglottis is a leaf-shaped cartilage that sits dorsal to the thryoid and superior to other laryngeal cartilages. The attached end (epiglottal stalk) is split in two. The alar is a horseshoe-shaped, rodent-specific structure that is located between the arytenoids and thyroid, and bounds the opening to the ventral pouch. In this species, the alar is connected to the epiglottal stalks. The connection points are on the ventral side of the alar cartilage near the rostral edge.

#### Ventral pouch

The ventral pouch is bounded by the thyroid cartilage ventrally and the alar cartilage at its dorsal entrance. The pouch is large, expanding medio-laterally and rostro-caudally forming a distinct, “w” shape (Figure 2 ventral pouch panel, top orientation).

### *In vitro* sound production mechanism

To identify whether singing mice use a MEAD (vocal membrane vibration) or aerodynamic whistle (wall impingement, edge-tone, cavity) mechanism (Figure 1D), we recorded sound and high-speed video of excised larynges (Figure 3A). All isolated larynges produced sound when the vocal folds (VFs) were adducted (5 males, 2 females, Table 1) and did not when VFs were not adducted (Table 1, “No Treatment”). High-speed imaging indicated that sound was produced in absence of tissue vibration (Video S1). These tones had a fundamental frequency between 22–46 kHz (µ = 33±3.5 kHz) which is almost entirely within the range of the singing mouse song (Figure 3D). Thus, we found support for whistle-based sound production in this species.

**Figure 3.**
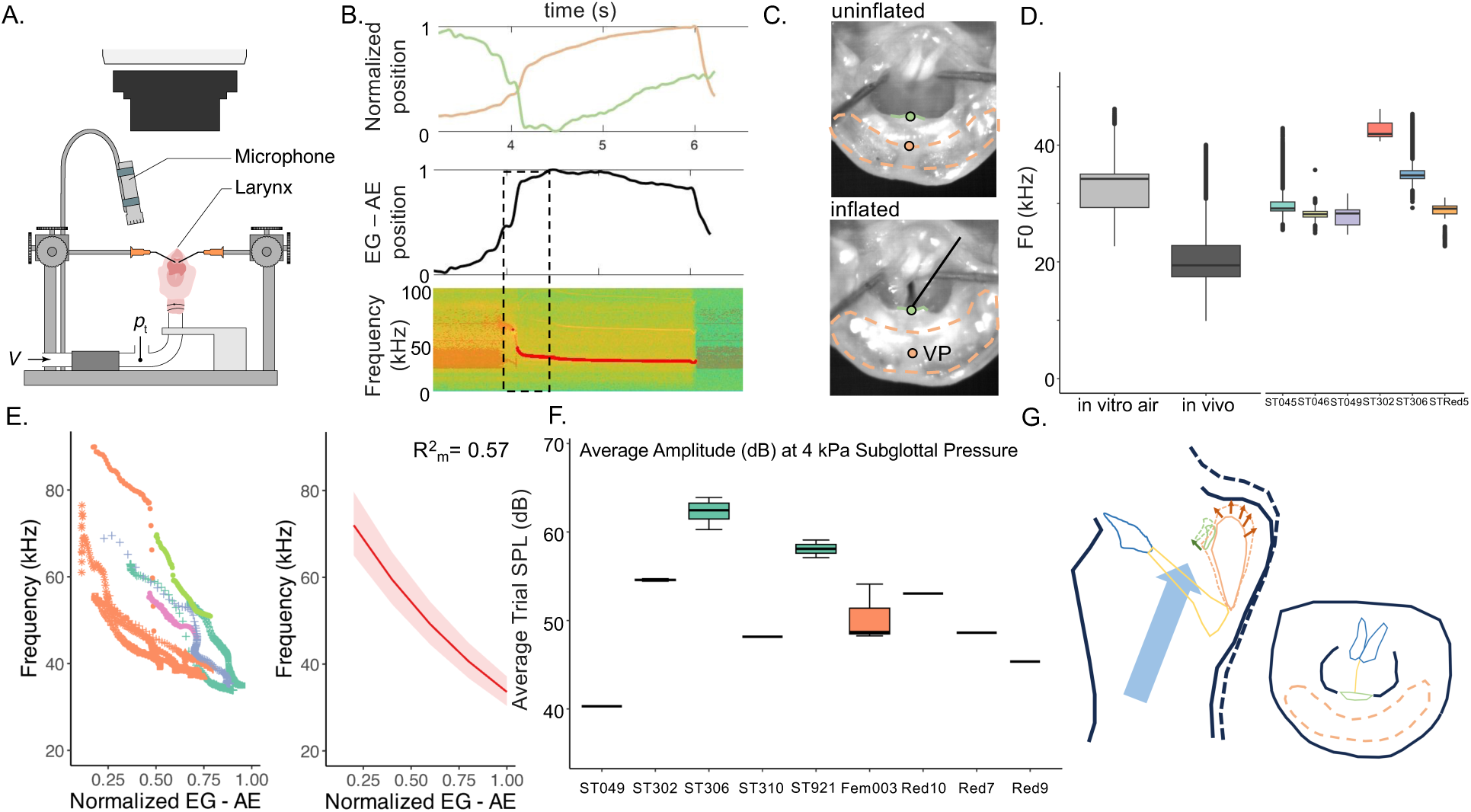
*In vitro* larynx manipulations suggest that singing mouse song is a whistle mechanism that requires ventral pouch inflation. **A.** Schematic of the experimental rig. B. (Top) Normalized landmark positions for alar edge (green) and tissue atop ventral pouch (located by epiglottal stems – EG, orange) across 1 adduction trial. (Middle) Difference between landmark positions across time. (Bottom) Sound spectrogram. Red line indicates F0 detected using Wiener’s entropy and the Yin algorithm. Black dotted box indicates time frame plotted in E. C. Still images from video showing uninflated (top) and inflated (bottom) alar pouch (see Video S1). Orange dashed lines indicate the ventral pouch. Green line indicates the alar edge. Orange and green dot represent landmark positions that are plotted in B. D. Fundamental frequency of sounds produced by mice *in vivo* (dark gray) and by excised larynges *in vitro* (light gray). Colored boxplots are frequencies produced by each larynx during adduction trials. E. (left) Relationship between landmark positions and frequency at sound onset. Color = larynx, dot type = trial. (right) Predicted values from mixed-effects model with 95% confidence interval (fixed effect: EG – AE position, random effect: larynx ID). F. Average SPL of sound for each larynx at 1 meter. Green represents larynges used in experiment 1. Orange represents larynges used in experiment 2. G. Sound production model. Mid-sagittal (left) slice and top-down (right) view of a larynx. Blue arrow represents air flow. Solid lines are laryngeal structures at rest. Dotted lines in top image represent the inflated position. Green and orange arrows show the direction of alar and ventral pouch movement respectively. V = velocity, p1 = air pressure, s = seconds, EG = epiglottis, AE = alar edge, VP = ventral pouch, F0 = fundamental frequency, In vitro air = whistle tones produced in excised larynges. In vivo = songs produced by mice in this study prior to euthanasia. Orange = ventral pouch. Green = alar, yellow = vocal folds, light blue = arytenoid.

**Table 1.** Number of larynges that produced whistles (“Tones”) or no sound in the following trials: no vocal fold adduction and an intact epiglottis cartilage (“No treatment”), vocal fold adduction and a cut epiglottis (“Adduction”), and vocal fold adduction, cut epiglottis, and cut alar (“Cut alar”). Whistles refer to sounds produced without tissue vibration (e.g., an edge-tone, wall impingement, or shallow cavity mechanism). “-“ = 0; M = male, F = female.

|  | # larynges (M,F) | Tones | No sound |
| --- | --- | --- | --- |
| No treatment | 8(6,2) | - | 8 |
| Adduction | 8(6,2) | 8 | - |
| Cut alar | 8(6,2) | - | 8 |

To distinguish between the potential whistle models (Fig 1D), we examined the structures necessary for sound production. We found that sound did not rely on an intact upper epiglottis, but did require an intact alar edge and ventral pouch (Table 1).

The singing mouse ventral pouch inflated during all adduction trials (Figure 3, Video S1). As air filled the pouch, the alar cartilage moved farther from the glottal opening, but also seemed to rotate as the edge was caught in the airflow. Inflation and alar movement began prior to sound production in all adduction trials. Across these trials, we used the distance between two landmarks (midway between epiglottal stalks = “EG” and the medial alar edge = “AE”) as a proxy for inflation (Figure 3B-C). The position of each of these landmarks and the distance between them were ∼50% of their maximal position at sound initiation (normalized EG = 0.459±0.248, normalized AE = 0.444±0.289, normalized difference = 0.491±0.258). We ran a linear mixed effects model on log_10_ transformed frequency (estimation method: maximum likelihood) and found a main effect of the normalized distance between EG and AE landmarks (β = - 0.41, SE = 0.008, Table S5, Figure 3E).

We next tested whether the alar cartilage was necessary for whistle production and found that cutting the superior portion of the alar (the top of the horseshoe see Figure 2) prevents sound production (Table 1). Because the alar cartilage forms the opening to the ventral pouch and connects to the soft tissue that forms the superior boundary of this air sac, destroying the alar cartilage also prevents pouch inflation.

An inflating pouch can alter frequency by changing the distance the air travels (“x” in Figure 1D) and by changing the volume of the cavity, if it acts as a resonator^11,14,49–51^. To understand whether sound can be produced without inflation, we filled the pouch with metal balls (5 larynges) or pieces of wax (3 larynges) (Table 2). In most trials, no sound or no stable frequencies were produced (19/21 trials) with little to no pouch inflation. In the three trials that produced substantial tones, there was an approximately 2-fold increase in frequency (Figure S3). Although we could not disentangle the contribution of the pouch in producing a stable whistle, together, these data indicate that the pouch and its inflation are essential to sound production.

**Table 2.** Number of pouch-fill trials that produced tones or no stable sound. F shift indicates whether the frequency was higher than frequencies produced by that larynx in the adduction trial. Columns separate trials by what was used to fill the pouch. F = frequency.

|  | metal balls | wax |
| --- | --- | --- |
| Tones |  |  |
| no F shift | 1 | - |
| F shift upwards | 1 | 1 |
| No stable sound | 11 | 7 |

The tones that were produced by an aerodynamic whistle with pouch inflation were loud. When a 4 kPa subglottal air pressure block was supplied, larynges produced a sound pressure level of ∼51 dB at 1 m (SD: 6.66; Figure 3F, Table S6).

Together, this evidence supports a whistle mechanism, likely an edge-tone or shallow cavity, that relies on an inflating ventral pouch to produce songs (Figure 3G).

### *In vitro* frequency control

Because the cricothyroid muscle is known to modulate frequency in both MEAD and whistle sounds in other species^3,37,52^, we mimicked the effect of CT contraction using a servo motor to rotate the cricoid cartilage *in vitro*. In these trials, we got whistles in 5 out of 6 larynges tested (4 M, 2 F) and tracked the midpoint of the thyroid’s superior edge in all videos. We found that upon cricoid rotation, the glottal opening narrowed (β = 3.17, SE = 0.055, Table S7, Video S2) leading to small increase in frequency (Figure 4, Table S8). This suggests that CT action can alter frequency by changing glottal opening width, which, alongside flow, contributes to air speed.

**Figure 4.**
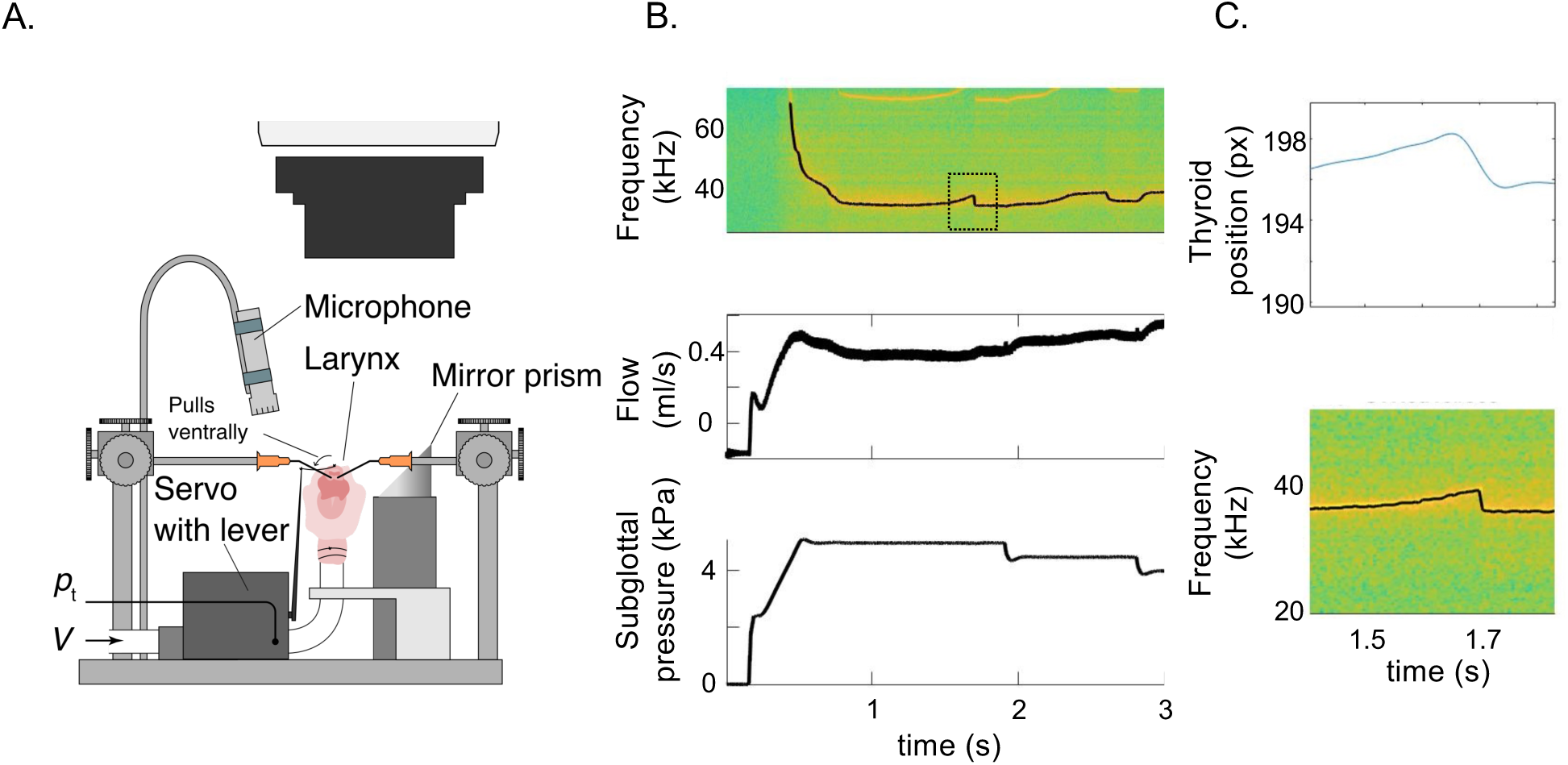
Imitating cricothyroid action *in vitro* increases frequency. A. Schematic of the experimental rig. B. Example larynx trial (STRed10) of frequency (top), air flow (middle), and subglottal pressure (bottom) across time. Black line on spectrogram (top) is frequency detected using Wiener’s entropy. Dotted gray box corresponds to inset in C. C. Pulling the thyroid cartilage increases sound frequency. s = seconds, px = pixels.

### Cricothyroid muscle ablation

To understand whether singing mice use the cricothyroid during singing, we recorded songs from mice where we surgically-ablated the CT muscle (see methods). The degree of ablation varied between the experimental animals, and we quantified the CT volume after song recording using histology.

Ablation animals had smaller estimated cricothyroid (CT) volumes (Welch’s t-test, *p* = 0.002) compared to control (sham-surgery) animals, but did not differ in the volumes of other laryngeal muscles (PCA: Welch’s t-test, *p* = 0.41; TA: Welch’s t-test, *p* = 0.90) (Figure S4).

While the ventral attachment points of the CT to the cricoid were fully ablated, the deeper (dorsal) and more lateral portions of the CT were less ablated in all individuals. Based on visual inspection of histological images, 2 animals had a minor CT ablation (D and E), 2 animals had a moderate CT ablation (B and C), and 1 animal had a major ablation (A). Surgery impacted other intrinsic laryngeal muscles in three individuals (A, C, E).

#### Qualitative song differences

All ablation individuals exhibited unique note phenotypes. Animals that had more complete cricothyroid ablations had greater reductions in frequency bandwidth (Figure 5A). The frequencies lost were variable at all levels from inter-individual to within a single song (see Figures S5-7 for examples). Some abnormal notes had a reduced frequency bandwidth where the upper, lower, or both ends of the frequency spectrum were lost (Figure 5A). Other notes lacked middle frequencies but retained normal frequency maxima and minima. Animals with moderate and severe ablations had note frequency reductions across the entire song, while animals with minor ablations showed frequency loss partway through the song, corresponding to where notes have the highest amplitudes and greatest frequency ranges.

**Figure 5.**
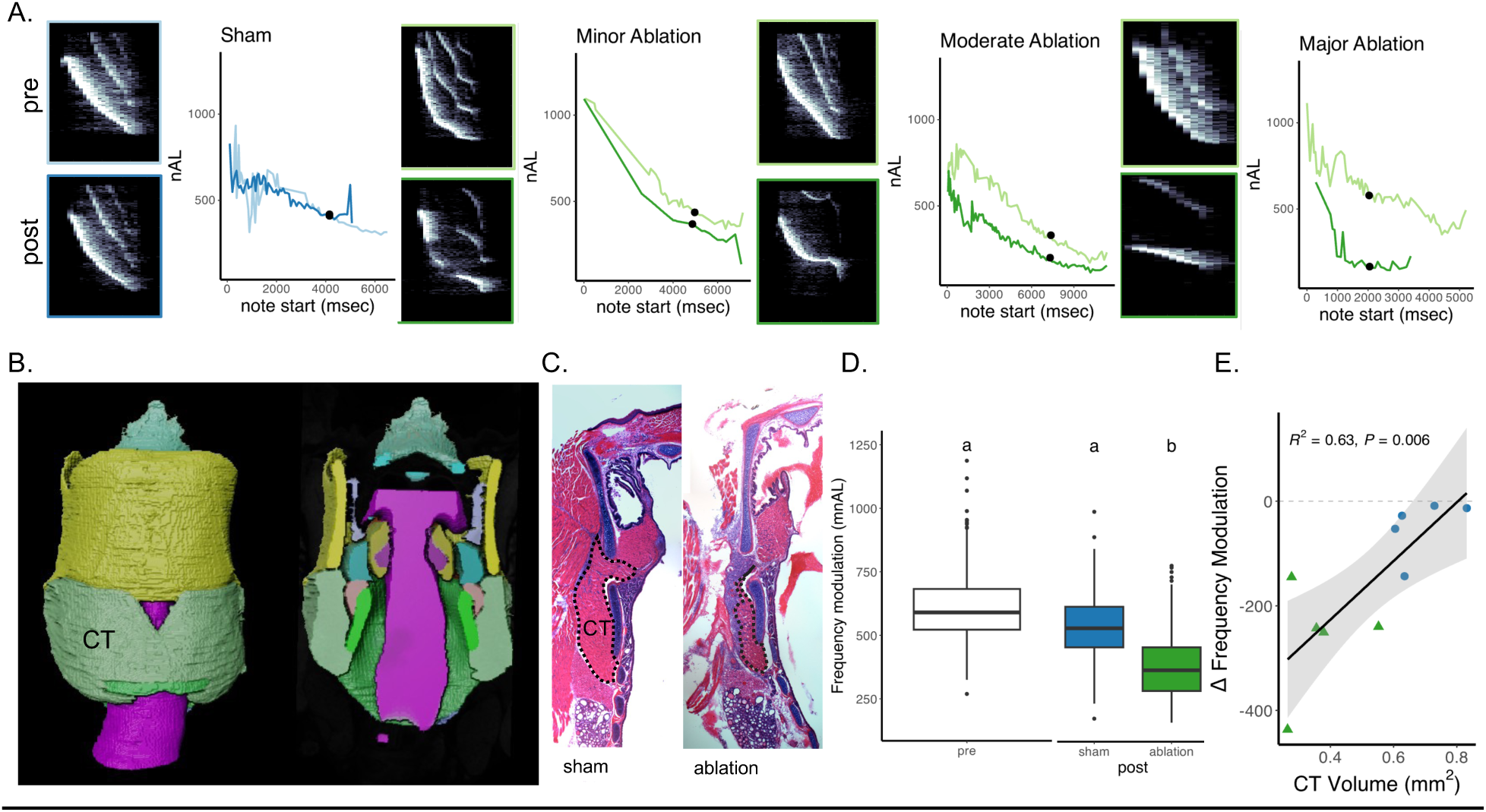
Cricothryoid (CT) muscle ablation impairs frequency modulation in singing mice. A. Location-matched exemplar notes from pre- and post-surgery songs. Each panel has a spectrogram from a pre- (top) and post-surgery (bottom) song and a plot showing the normalized arc length across each song. The point on each graph shows the location of the notes in the spectrograms. B. Exemplar μCT reconstruction of the cartilages, muscles, and airway of a singing mouse larynx. C. H&E-stained, transverse section of a sham (left) and ablation (right) larynx. Dotted line outlines the intact CT muscle (pink). D. Mean, normalized arc lengths are reduced in post-surgery songs of ablation animals compared to all other groups (dark green). E. Ablation individuals have smaller CT volumes and larger differences in mean, nAL between timepoints (post-pre mnAL). CT volume was calculated from H&E-stained laryngeal sections. Only intact muscle cells were measured. Light colors = pre-surgery, dark colors = post-surgery. Blue – sham, green = ablation. CT = cricothyroid.

#### Quantitative song differences

Mean, normalized arc length (mnAL, Figure 1C) was lower in the post-surgery period (Wilcoxon test, 5/5 ablation *p*<0.001; 2/5 sham *p*<0.001; 1/5 sham *p*<0.05, 2/5 sham *ns*, Figure S8). To understand the impact of CT ablation on song, we fit linear mixed-effects models for mnAL. Likelihood ratio tests and an AIC model selection approach indicated that the best model was a random slopes model which included timepoint, treatment, and their interaction as fixed effects and individual identity as a random effect (Table 3). We used a random slopes model to account for differences in surgery across subjects. This model indicated that post-surgery ablation songs have notes with smaller arc lengths compared to all other songs (Figure 5D). One mouse (A) had a CT ablation which impacted other intrinsic muscles. We ran the same linear mixed-effects models on the dataset excluding individual A and found that the best model contained the same terms as the full dataset model with similar coefficients (Table S9).

**Table 3.** Coefficients, confidence intervals (CI), p-values, and R^2^ of the model that best explains the variance in mean normalized arc length (mnAL).

| Predictors | Estimates | CI | *p* |
| --- | --- | --- | --- |
| (Intercept) | 600.21 | 542.94 – 657.48 | **<0.001** |
| Treatment [ablation] | 18.28 | -62.96 – 99.53 | 0.659 |
| Timepoint [post] | -49.95 | -114.53 – 14.64 | 0.130 |
| Treatment [ablation] * Timepoint [post] | -209.30 | -301.52 - -117.09 | **<0.001** |
| Marginal R<sup>2</sup> / Conditional R<sup>2</sup> | 0.358 / 0.613 |  |  |

We used AIC model selection to find the combination of muscle volumes that best predict the difference in mean, normalized arc length across timepoints (Figure 5E). Estimated CT volume alone was the best model (*p* < 0.005, R^2^ = 0.63) and was responsible for 77% of the cumulative model weight (Table 4). In summary, singing mice with cricothyroid muscles that were surgically reduced in volume produced songs with smaller frequency bandwidths, indicating that this muscle is necessary for normal frequency modulation.

**Table 4.** Model selection results for mnAL ∼ muscle volume. K = number of parameters. AICc = AIC value of the model corrected for a small sample size. Delta AICc = difference between the AIC value of the best model compared to the model being examined. AICc Wt = proportion of total predictive power. Cumul Wt = cumulative sum of AIC weights. LL = log-likelihood. CT = cricothyroid, LCA = lateral cricoarytenoid, TA = thyroarytenoid.

| | K | AICc | ΔAICc | AICc Wt | Cumul. Wt | LL |
| --- | --- | --- | --- | --- | --- | --- |
| CT | 3 | 125.92 | 0.00 | 0.77 | 0.77 | -57.96 |
| CT.LCA | 4 | 130.75 | 4.82 | 0.07 | 0.84 | -57.37 |
| LCA | 3 | 130.98 | 5.06 | 0.06 | 0.90 | -60.49 |
| null | 2 | 131.61 | 5.68 | 0.05 | 0.95 | -62.95 |
| CT.TA | 4 | 131.92 | 5.99 | 0.04 | 0.99 | -57.96 |
| TA | 3 | 134.46 | 8.54 | 0.01 | 1.0 | -62.23 |
| CT.LCA.TA | 5 | 136.84 | 10.92 | 0.00 | 1.0 | -55.92 |

## Discussion

Rodents use unique morphology and multiple sound production mechanisms to produce their diverse vocalizations^10,14,53^. We examined the mechanisms and morphology underlying the long-distance songs of Alston’s singing mouse, *Scotinomys teguina*^12,52^. We considered vocal membrane vibration, as seen in bats^54^, and aerodynamic whistles, as exhibited by some rodents^10,11,14,18,19^, because these mechanisms can produce sounds within the frequency range of the singing mouse song. Using *in vitro* and *in vivo* methods, we found that singing mice use a novel aerodynamic whistle mechanism that relies on the inflation of a large ventral pouch. Our assays indicated that air flow and the cricothyroid likely control song frequency. We are the first to report airflow driven ventral pouch inflation as an essential facet of whistle sound production. Below, we discuss our working model of sound production and frequency control as well as its implications for the study of vocal mechanisms.

### Sound production mechanism

Our *in vitro* experiments indicated that singing mice use a whistle mechanism that is unique because it requires ventral pouch inflation to produce sound. During experimentation, we considered both aerodynamic whistles^10,11,14,17,55^ and myoelastic-aerodynamic (MEAD^15^) mechanisms since both can facilitate high-frequency vocalizations in rodents^12,18,19^. Although singing mice have a vocal membrane^48^, we found no evidence that the vocal membrane vibrates in the frequencies of the song.

Instead, we found that tones within the song frequency range were produced by an aerodynamic whistle that required an intact alar cartilage and inflating pouch. Ventral pouch inflation occurred in all trials that produced sound and began prior to sound initiation. We had to supply high subglottal air pressures at the beginning of trials to produce sound. Blocks of high pressure and decreasing ramps induced rapid pouch inflation and produced clear sounds, while ramps of increasing pressure produced slow and inconsistent pouch inflation and limited sound. In trials where we cut the alar edge, the ventral pouch was also damaged, as this cartilage bounds the entrance to the ventral pouch, and no sounds were made. To disentangle the effect of the alar edge and inflating ventral pouch on sound production, we filled this air sac. Preventing inflation by filling the pouch also inhibited sound in nearly all trials, suggesting inflation is essential to generate sound. Although we know that ventral pouch morphology varies among rodents^11,13^ and it has been suggested that this structure contributes to vocal differences, we are the first to find evidence that ventral pouch inflation plays a role in the production of sound.

The reliance on the alar cartilage and ventral pouch to produce sound indicates that song is likely produced by an edge-tone^11,55^ or shallow cavity^14^ mechanism. By filling the ventral pouch, we prevented air circulation within the sac; this interrupts shear layer formation and eliminates sounds produced by a shallow cavity. This perturbation does not disturb the alar edge, and so should in theory not alter edge-tone sounds^11,14,55^. However, we found ventral pouch inflation altered the position of the alar edge, which offers an unexpected mechanism by which blocking inflation could also prevent an edge tone. Our data cannot yet distinguish between an edge-tone or shallow cavity mechanism in singing mice.

In addition to its role in generating sound, it has been suggested that a dynamically changing ventral pouch may modulate frequency via its resonant properties^11^. Although most of our pouch fill trials produced no sound, a small subset of these trials (2/21) generated tones of ∼2x higher frequency. These results could be consistent with a scenario where the ventral pouch acts as a resonator whose volume has decreased. Mathematical and mechanical models based on singing mouse laryngeal anatomy would be feasible next steps towards pinpointing which feedback mechanisms causes the whistle to stabilize^14^ (i.e., alar-edge or shallow cavity), as well as investigating the resonant properties of the ventral pouch. Nevertheless, higher subglottal pressures alongside the relatively low frequency range of this whistle, likely enables the production of an omnidirectional, long-distance, advertisement signal.

Although high subglottal air pressures facilitate whistles in this species, the singing mouse song is quieter than the long-distance calls of other Neotomids. For example, compared to the estimated 5 m active space of singing mouse songs^22,26^ (∼60 dB SPL at 1 m), *Onychomys* produce calls with 85.7 dB SPL at 1 m (50 m active space^56^) and *Peromyscus truei* advertisement calls have 72.1 dB SPL (12.4 m active space^57^). Unlike singing mouse songs, *Onychomys*, *Peromyscus*, and close relatives in *Reithrodontomys* use tissue vibration to produce their long-distance calls. This may contribute to the differences in amplitude across these species.

The relationship between the elaboration of pouch morphology and vocalization across rodents suggests that pouch shape and size contribute to species differences in vocal features. Laboratory rats and mice (family: Muridae) have small, ovate ventral pouches^11^ which do not play a major role in the production of their short, highly-directional, quiet, high-frequency vocalizations (USVs)^5,14^. Non-Baiomyini Neotomids (*Onychomys*^12^*, Reithrodontomys*^58^*, Peromyscus*^18^) use whistle mechanisms to produce close-range USVs and have ovoid pouches like lab rodents. Kangaroo rats (family: Heteromyidae, genus: *Dipodomys*) lack a ventral pouch and alar cartilage^11^ and have no evidence of USV production, although they make a limited repertoire of other vocalizations^59–61^. Conversely, pygmy mice (genus: *Baiomys*) use a whistle mechanism for both close- and long-range communication and have large, “w”-shaped pouches^13^. Like their close relative, singing mice have large “w”-shaped pouches^13^ and use a whistle mechanism for their long-distance songs. Compared to the whistles of other rodents, singing mouse songs are incredibly lengthy and stereotyped, suggesting unique features of the singing mouse sound production mechanism, like an enlarged and inflating ventral pouch may facilitate their precise physical control or enable particular features of song.

Laryngeal air sacs are not limited to rodents but can be found among primates^37,62–65^, cetaceans^66^, pinnipeds^67^, ungulates^68^, carnivores^69^, chiroptera^70^, and marsupials^71^. They vary in their location, shape, and size, as well as by whether they can inflate or are bounded by bony structures (e.g., some subhyoidal sacs), which presumably impacts their acoustic functions^62^. Laryngeal air sacs might alter MEAD-based vocalizations by amplifying sound, filtering frequencies, or making sound production more energetically efficient^64^, but few of these hypotheses have been tested. Our results suggest a new role for air sacs in aerodynamic whistles: their inflation is essential to generate sound. More research is needed to understand how air sacs impact MEAD-based and whistle sounds. Connecting morphology to their underlying function will allow us to begin to explore how variation in morphology and biomechanics contributes to the evolution of novel vocalizations.

### Mechanisms of frequency modulation

Having identified that *Scotinomys* songs are aerodynamic whistles, we next asked how such vocalizations might be modulated. In laryngeal myoelastic-aerodynamic (MEAD) mechanisms, the cricothyroid muscle (CT) controls frequency. Its contraction increases frequency by increasing tension of the vibrating tissues^15^. For aerodynamic whistles, a computational model of frequency control in rats showed that any muscle that alters air speed or the distance that air travels might alter frequency^14^ (Fig. 1D). This includes the respiratory muscles that shape subglottal air pressure, and the CT and thyroarytenoid (TA) muscles^52^. The model indicated that TA action primarily affects whistle stability while CT action impacts whistle stability and decreases frequency in the rat larynx^14^.

Unlike rats, CT action appears to increase frequency in the singing mouse larynx. Whether CT action increases or decreases frequency depends on its dominant effect on laryngeal geometry. In rats, the main effect of CT action is to increase the distance between the glottal opening and alar edge. Instead of increasing the distance air travels and thereby reducing frequency, the dominant effect in our trials was glottal narrowing, which increases airspeed and therefore frequency. However, this outcome may differ if subglottal pressure changes concurrently with CT action. A computational model driven by muscle activity during singing could further clarify the role of the CT and other muscles.

Although our *in vitro* work indicated that CT contraction changed whistle frequency, we wanted to test whether singing mice use this muscle *in vivo*. We found that damaging the CT muscle reduces song frequency bandwidth. Damaging muscle fibers induces contractions with weaker forces, which produce smaller rotations of the thyroid cartilage, altering glottal opening size and the glottis-to-alar distance less dramatically. Although frequency bandwidth reduction was consistent, the exact frequencies lost varied between and within individuals and even within a single song. This is unusual, as singing mouse songs are highly stereotyped^29^. Some of the intraindividual variation in frequency range may be due to compensatory mechanisms. Hearing the abnormalities in their own songs, singing mice might use respiratory and other laryngeal muscles to attempt to recover their natural frequency range. Measuring respiratory and laryngeal muscle action post-surgery would shed light on this possibility.

Although the songs of singing mice and the USVs of laboratory mice and rats are both produced using whistle mechanisms, they differ greatly in frequency bandwidth and modulation. Singing mouse songs are highly stereotyped, particularly repeatable in frequency related measures. USV syllables of laboratory rodents are highly variable, although they can be organized into types by their shape and frequency^5^. Although there is a lot of frequency modulation in laboratory rodent USVs, not all of it is controlled actively by respiratory and laryngeal muscles. For example, the whistle model proposed by Håkansson et al.^14^ suggests that the frequency jumps used to characterize syllable types are stable whistle modes and are not induced by muscle action. Singing mouse songs do not feature these elements. Each note consists of a frequency downsweep across ∼30 kHz and we find that damaging one of their vocal muscles interrupted their ability to precisely control this song feature. Thus, frequency seems to be precisely and actively controlled in singing mice. Because of this, muscles like the cricothyroid might play an outsized role in the vocalizations of singing mice.

### Conclusions

Using *in vitro* and *in vivo* procedures, we find evidence that singing mice use a whistle mechanism that relies on an inflating ventral pouch and stabilized alar cartilage to produce their long-distance song. Our results indicate that an aerodynamic whistle can produce loud and precisely frequency-modulated advertisement vocalizations. Although we are not able to distinguish between an edge- and cavity model with these experiments, both models posit that the alar cartilage and ventral pouch are essential for sound production. They also tell us that laryngeal muscles that control air speed or distance, like the cricothyroid, are candidates for controlling sound frequency. Ventral pouch inflation contributing to the generation of stable acoustic waves is unique to singing mouse sound production and suggests a novel role for air sacs in mammalian sound production. Further work is warranted to understand how morphological variation may contribute to differences in acoustic structure across species and the ventral pouch seems a particularly promising candidate, since its morphology varies across species and is absent in at least one minimally-vocalizing rodent^11,13^.

Overall, our results add to mounting evidence that changes to both vocal anatomy (e.g., ventral pouch morphology) and biomechanics (e.g., whistle vs. MEAD mechanisms) are responsible for differences in vocalizations across rodents. Although the aerodynamic whistles that muroid rodents rely on to produce high frequency sounds are unique to this taxonomic group, the use of an air sac to shape novel sounds is part of a recurring theme that has evolved repeatedly across terrestrial vertebrates^63,64,72–76^. Our examination of the anatomy and biomechanics of the singing mouse larynx is a step towards understanding how novel vocalizations arise.

### Ethics

All animal procedures were approved by the University of Texas at Austin and New York University Grossman School of Medicine IACUC. All methods were done in accordance with IACUC regulations. We confirm that we have reported this research following the ARRIVE guidelines for the reporting of animal research.

### Data, Code, and Materials

Micro-CT data were deposited on the Texas Data Repository (https://dataverse.tdl.org/dataverse/uCTsteg). Excised larynx data and code were deposited on Github (https://github.com/samksmith/singing-mouse-SPM) and the TDR (https://dataverse.tdl.org/dataverse/STeg_SPM). The muscle ablation data and code were deposited on Github (https://github.com/samksmith/steg_muscle_ablation). The histological data associated with the muscle ablation study were deposited on figshare [Project: Cricothyroid Muscle Ablation in Alston’s singing mouse, (Scotinomys teguina)].

## Supporting information

Main Supplementals

## Acknowledgements

We would like to thank Dr. Jessie Maisano and the University of Texas High-Resolution X-Ray CT Facility (UTCT) for supporting the microCT data presented in this paper and for providing training on data visualization and analysis. UTCT is supported by the NSF Division of Earth Science Instrumentation and Facilities Program (NSF EAR-1762458), and NASA (80NSSC23K0199). Larynx sectioning for the muscle ablation study was done at the Center for Biomedical Research Support Microscopy and Imaging Facility at UT Austin (RRID:SCR021756). We thank members of the Phelps and Elemans laboratories for feedback on early conceptualizations, analysis, and visualization of results of this project.

## Authors’ contributions

Conceptualization – SKS, SMP, CE

Data curation – SKS, JH, PWF

Formal analysis – SKS, JH

Funding acquisition – SMP, ML

Investigation – SKS, JH, PWF

Methodology – SKS, CE, JH, SMP

Project administration – SKS, SMP, CE

Software – SKS, JH, SMP, CE

Resources – SMP, CE, ML

Supervision – SMP, CE, ML

Validation – SKS, JH, PWF

Visualization – SKS, JH

Writing – original draft – SKS

Writing – review & editing – SKS, SMP, CE, JH, ML, PF

