## Supplementary material for "Mechanisms and control of a novel vocalization: The singing mouse song is a whistle that depends on air sac inflation": Main Supplementals

| Image reconstruction parameters | |
|---|---|
| center shift | -2.5 |
| beam hardening | 0.6 |
| theta | -10 |
| byte scaling | [-100, 3200] |
| binning | 1 |
| recon filter smoothing kernel size | 0.5 |

**Table S1.** Image reconstruction parameters for all larynx scans (n=6, 3 males, 3 females).

| Sample | Species | Sex | Total Slices | XYZ |
|---|---|---|---|---|
| 978A | *S. teguina* | M | 894 | [-50, 14194, -1] |
| 981A | *S. teguina* | M | 970 | [-219, 23873, 218] |
| 982A | *S. teguina* | M | 919 | [529, 18828, -83] |
| 990A | *S. teguina* | F | 995 | [-50, 14194, -1] |
| 991A | *S. teguina* | F | 984 | [219, 23873, 218] |
| 992A | *S. teguina* | F | 978 | [3, 23049, -34] |

**Table S2.** Total number of slices, and XYZ values for each µCT sample.

| Structure | Number of Landmarks |
|---|---|
| Left glottal edge | 6 |
| Right glottal edge | 6 |
| Alar edge | 5 |
| Epiglottis | 5 |

**Table S3.** Landmarks placed on high-speed larynx videos.

### Deep learning algorithm for analyzing video data

We used DeepLabCut[1], a software package that facilitates markerless pose estimation using deep neural networks that have been supplied a relatively small amount of training data. We trained a different DeepLabCut network per larynx in the dataset using DeepLabCut v. 2.3.0 (single animal network). We selected videos where the larynx moved in different ways to generate a good training dataset to capture all the ways the larynx can moved in our experiment. We used kmeans clustering to select 15 frames per video. This procedure downsamples the video and clusters frames using k-means. Frames are then selected from different clusters to ensure that the frames selected look different from one another. Frames were manually labeled using the DLC GUI. For experiment 1, 6 landmarks were placed on the left edge of the glottal opening, 6 on the right side of the glottal opening, 5 placed along the alar edge, 4 placed around the arytenoid stalks, and 4 placed along the thyroid edge (**Figure S1**).

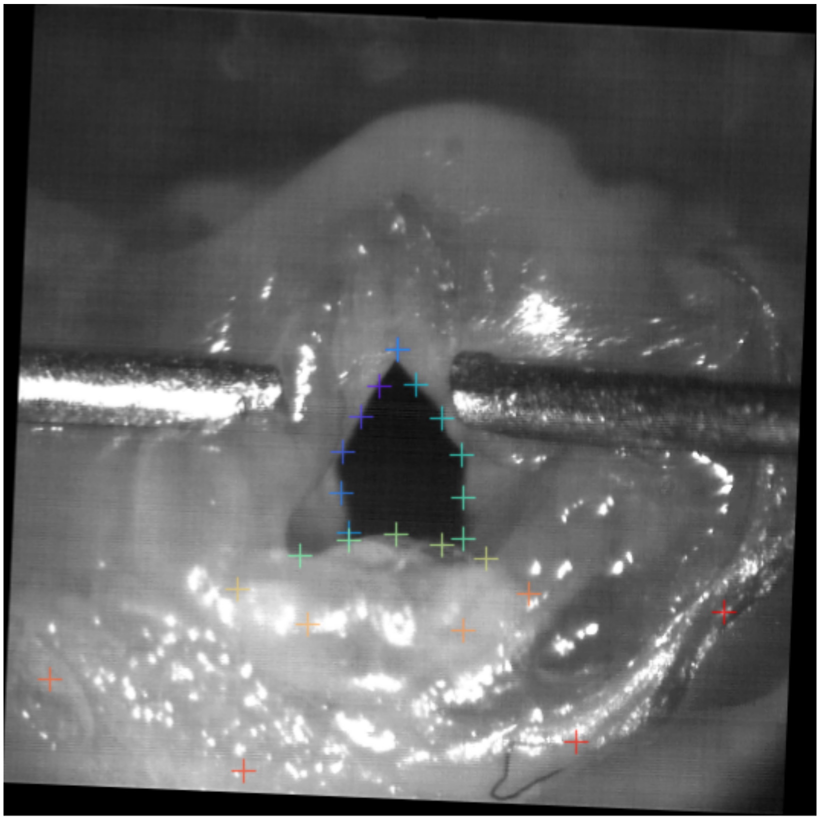

**Figure S1.** An example labeled frame from a video used to train a larynx from experiment 1. Colored plus signs are landmarks. Dark blue/purple landmarks are placed along the left glottal edge, dark blue/green landmarks denote the right glottal edge, the light green/yellow landmarks were placed along the alar edge, light orange landmarks were placed around the epiglottal stalks, and the red landmarks were placed along the thyroid edge.

We then generated the training dataset. This is done by combining the labeled datasets from all the videos for a single larynx and splitting them to create training datasets that train the network and testing datasets which are used to evaluate the network. Videos were not downscaled for training (global scale = 1.0). We used a batch size of 2 and initial weights resnet_v1_50. Other than these parameters, we used default settings for a single animal network. We trained each network for up to 600,000 iterations, with a snapshot saved every 50,000 iterations. After training, each snapshot was evaluated using the built-in DLC function for evaluating networks and the iterations that corresponded to the snapshot with the lowest filtered pixel error was selected to use for analyzing the videos. The parameters and other metadata from the training for each larynx are included below in Table SX. The resulting coordinates were low-pass filtered at 10 Hz to reduce the jitter artifacts due to the relatively low video resolution.

| Experiment | Sample | # of videos trained on | Total frames | Training iterations | p_cut_test_error | Max iterations |
|---|---|---|---|---|---|---|
| 1 | STRed5-SamKSmith-2023-09-13 | 4 | 60 | 600000 | 3.32 | 600000 |
| 1 | ST_production_mech_046-Samantha K. Smith-2023-03-06 | 4 | 60 | 450000 | 2.09 | 500000 |
| 1 | ST_production_mech_049-SamanthaKSmith-2023-03-24 | 4 | 60 | 450000 | 1.64 | 500000 |
| 1 | ST045#009-SamKSmith-2023-09-22 | 1 | 15 | 600000 | 0.63 | 600000 |
| 1 | ST045-SamKSmith-2023-09-13 | 3 | 45 | 550000 | 1.97 | 600000 |
| 1 | ST302#007-SamKSmith-2023-09-22 | 1 | 15 | 350000 | 1.66 | 600000 |
| 1 | ST302-SamKSmith-2023-09-12 | 4 | 60 | 250000 | 1.95 | 600000 |
| 1 | ST306-SamKSmith-2023-09-12 | 4 | 60 | 600000 | 1.68 | 600000 |
| 1 | ST901-SamKSmith-2023-09-13 | 3 | 45 | 400000 | 1.46 | 600000 |
| 1 | STRed5#006-SamKSmith-2023-10-03 | 1 | 15 | 250000 | 2.09 | 600000 |

| | | | | | | |
|---|---|---|---|---|---|---|
| 2 | STRed10-SamKSmith-2023-09-16 | 2 | 30 | 250000 | 1.97 | 600000 |
| 2 | ST310-SamKSmith-2023-09-17 | 3 | 45 | 500000 | 1.59 | 600000 |
| 2 | STF003-SamKSmith-2023-09-13 | 2 | 30 | 300000 | 1.98 | 600000 |
| 2 | STF891-SamKSmith-2023-09-17 | 3 | 45 | 600000 | 2.71 | 600000 |
| 2 | STRed7-SamKSmith-2023-09-14 | 2 | 30 | 350000 | 1.8 | 600000 |
| 2 | STRed9-SamKSmith-2023-09-17 | 2 | 30 | 300000 | 1.66 | 600000 |

**Table S4.** Metadata on the training of a deep learning algorithm on larynx videos.

We looked at the resulting labeled videos and identified a few that were not labeled accurately. For these videos, we trained a separate network (for example video #6 of STRed5). For a few of the videos, we accidentally chose snapshots that did not have the lowest pixel error. However, in each of these cases, the number of training iterations chosen did not massively exceed the lowest pixel error value. For example, for larynx 310, 300,000 iterations was best with a pixel error of 2.52, but we chose 600,000 iterations with an error of 2.71. Because we verified the resulting coordinates by generating and viewing labeled videos, we feel confident that our position data was accurate enough to facilitate downstream analyses.

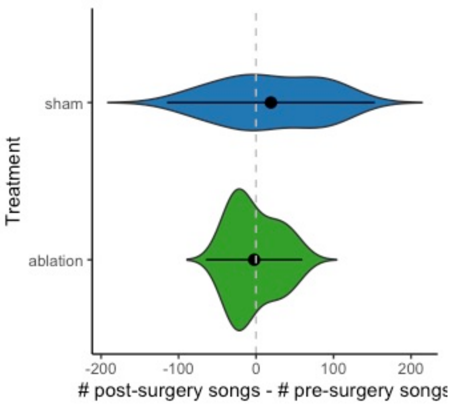


**Figure S2.** No differences in number of songs sang at the two time points for sham or ablation animals.

| | **log10(freq/1000)** | | |
|---|---|---|---|
| *Predictors* | *Estimates* | *CI* | *p* |
| (Intercept) | 1.94 | 1.90 – 1.98 | **<0.001** |
| diffnorm | -0.41 | -0.43 – -0.40 | **<0.001** |
| **Random Effects** | | | |
| $\sigma^2$ | 0.00 | | |
| $\tau_{00\ id}$ | 0.00 | | |
| ICC | 0.54 | | |
| $N_{id}$ | 5 | | |
| Observations | 1386 | | |
| Marginal $R^2$ / Conditional $R^2$ | 0.574 / 0.802 | | |

**Table S5.** Results of mixed-effects model for $\log_{10}$ transformed frequency (kHz). Diffnorm = Normalized difference between EG and AE landmark positions.

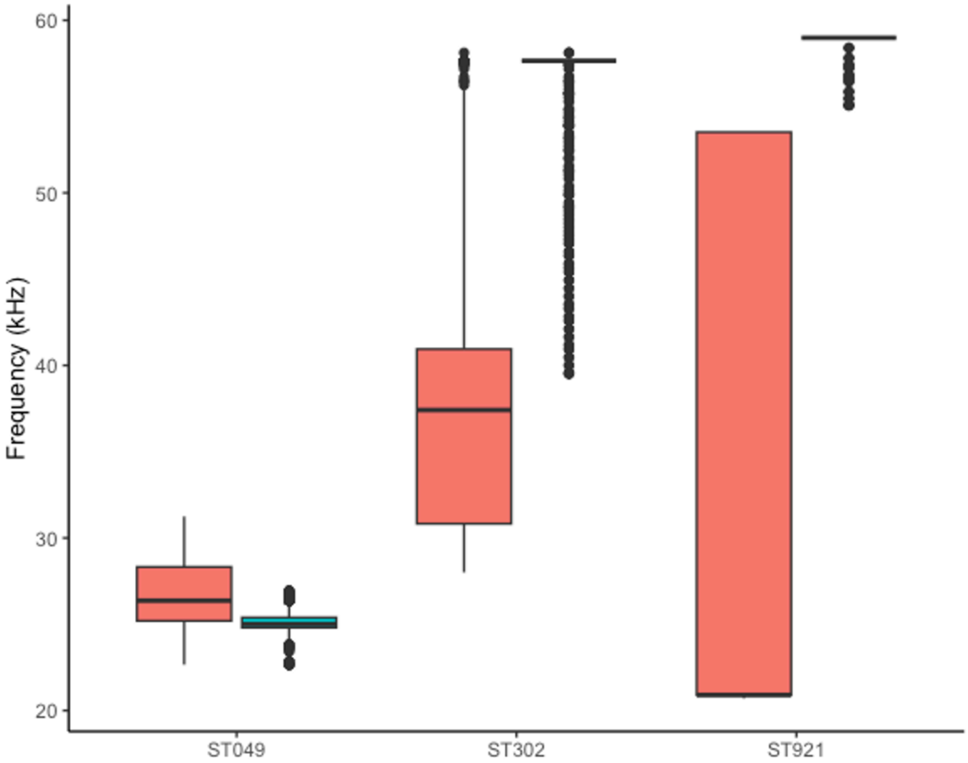


**Figure S3.** Boxplots of frequencies detected using Shannon’s entropy in adduction trials (pink) and pouch fill trials (blue). Only individuals and trials shown where substantial sound was produced upon pouch filling. The five other larynges produced little to no sound when the pouch was filled with metal balls or wax.

| Larynx ID | Experiment | Mean source level (dB re. 20μPa at 1 m) | SD |
|---|---|---|---|
| ST049 | 1 | 40.3 | - |
| ST302 | 1 | 54.6 | 0.2 |
| ST306 | 1 | 62.3 | 1.57 |
| ST310 | 2 | 48.2 | - |
| ST921 | 1 | 58.1 | 1.41 |
| STFem003 | 2 | 50.3 | 3.26 |
| STRed10 | 2 | 53.0 | - |
| STRed7 | 2 | 48.6 | - |
| STRed9 | 2 | 45.4 | - |

**Table S6.** Mean and standard deviation (SD) of source level (dB relative to 20μPa at 1 m) for each larynx from trials where subglottal air pressure was 4 kPa. Experiment 1 refers to the sound production mechanism experiment. Experiment 2 refers to the physical control experiment. dB is reported at 1 m distance assuming transmission loss according to spherical spreading. Microphone distance in original experiment was 3.9 cm.

| | GA | | |
|---|---|---|---|
| *Predictors* | *Estimates* | *CI* | *p* |
| (Intercept) | -488.04 | -555.59 – -420.49 | **<0.001** |
| Thpt | 3.17 | 3.06 – 3.28 | **<0.001** |
| **Random Effects** | | | |
| $\sigma^2$ | 8899.53 | | |
| $\tau_{00\ id}$ | 4994.94 | | |
| ICC | 0.36 | | |
| $N_{id}$ | 5 | | |
| Observations | 10353 | | |
| Marginal $R^2$ / Conditional $R^2$ | 0.465 / 0.657 | | |

**Table S7.** Results of mixed-effects model for glottal area (GA). Thpt = position of thyroid landmark in pixels.

| | log10(freq/1000) | | |
|---|---|---|---|
| *Predictors* | *Estimates* | *CI* | *p* |
| (Intercept) | 1.55 | 1.47 – 1.63 | **<0.001** |
| flowml | -0.05 | -0.06 – -0.05 | **<0.001** |
| Thpt | 0.00 | 0.00 – 0.00 | **<0.001** |
| GA | -0.00 | -0.00 – -0.00 | **<0.001** |
| **Random Effects** | | | |
| $\sigma^2$ | 0.00 | | |
| $\tau_{00\ id}$ | 0.01 | | |
| ICC | 0.82 | | |
| $N_{id}$ | 5 | | |
| Observations | 10353 | | |
| Marginal $R^2$ / Conditional $R^2$ | 0.198 / 0.857 | | |

**Table S8.** Results of mixed-effects model for $log_{10}$-transformed frequency (kHz). Flowml = air flow in milliliters. Thpt = position of thyroid landmark in pixels. GA = glottal area in pixels$^2$.

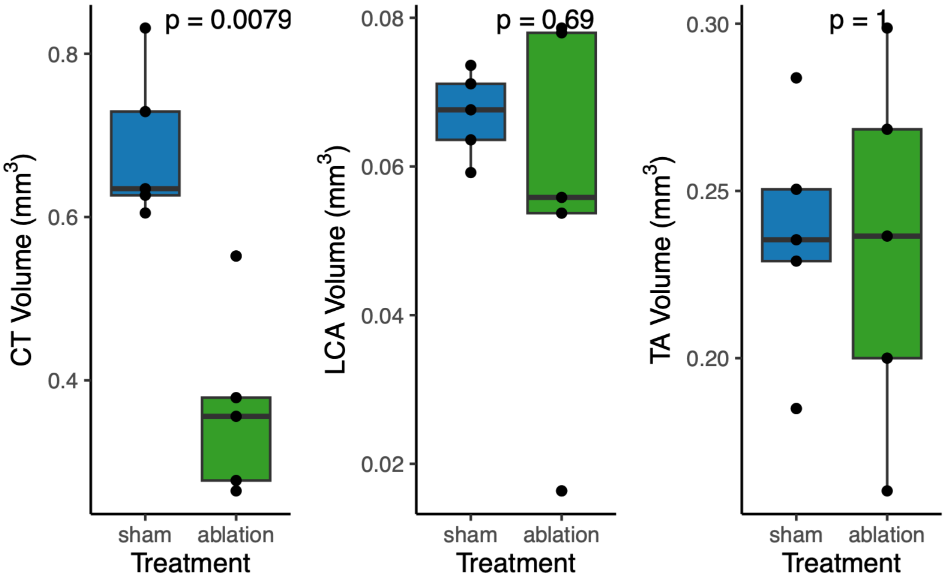


**Figure S4.** Ablation animals had smaller CT volumes than sham animals. All other measured muscle volumes did not differ between the two groups.

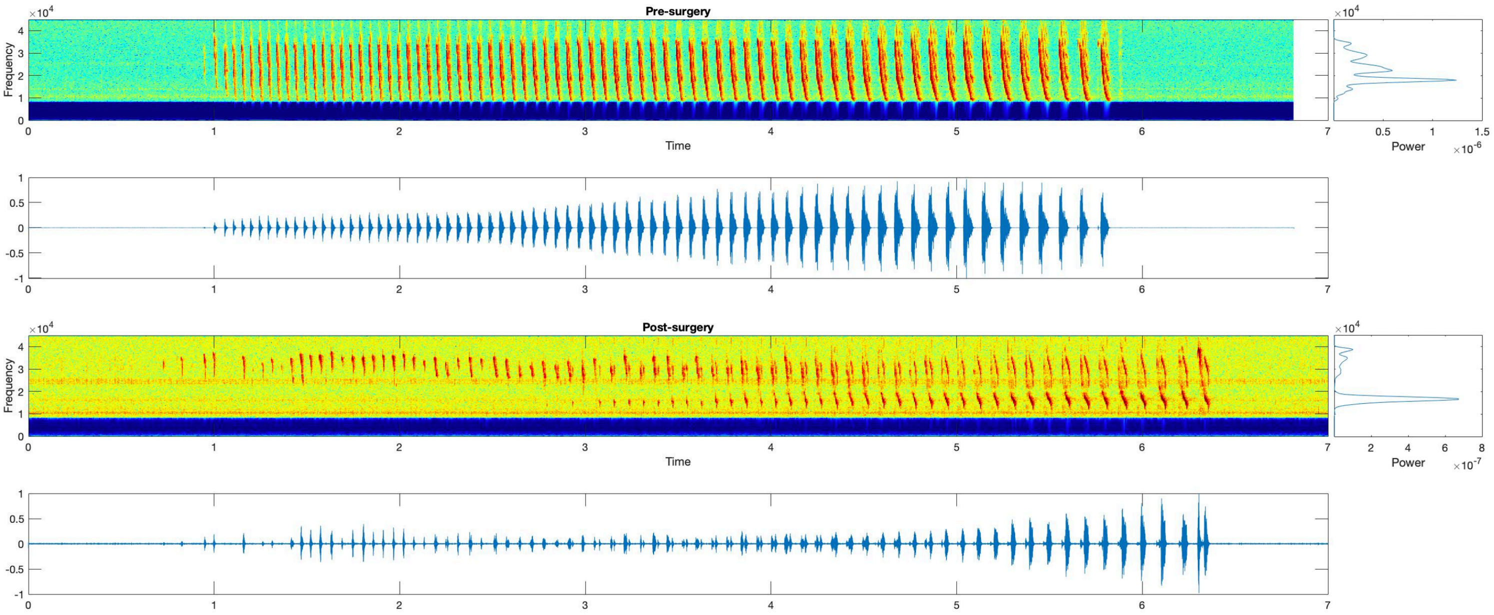


**Figure S5.** Spectrograms and oscillograms of a pre- (top) and post-surgery (bottom) song for an animal that had a severe ablation (Individual A). Post-surgery song has a severely reduced bandwidth, losing both upper and lower ends of the frequency range.

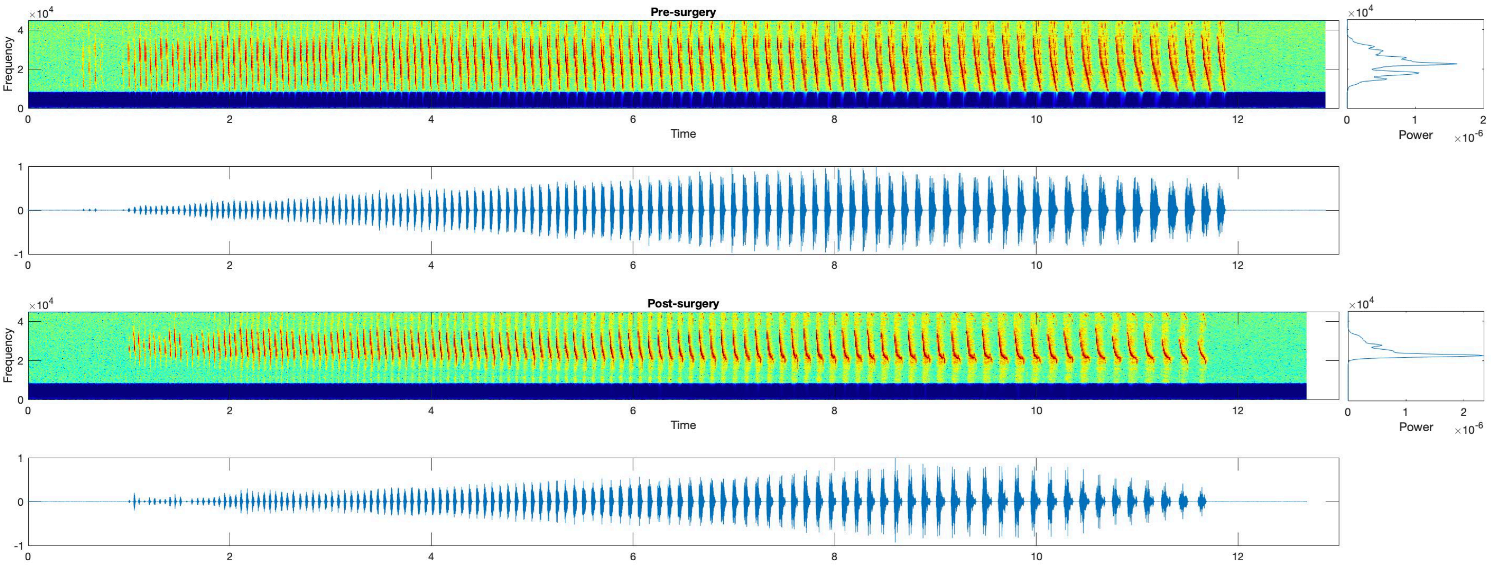


**Figure S6.** Spectrograms and oscillograms of a pre- (top) and post-surgery (bottom) song for an animal that had a moderate ablation (individual B). Post-surgery song has a reduced frequency bandwidth, where frequencies below 20 kHz have been lost.

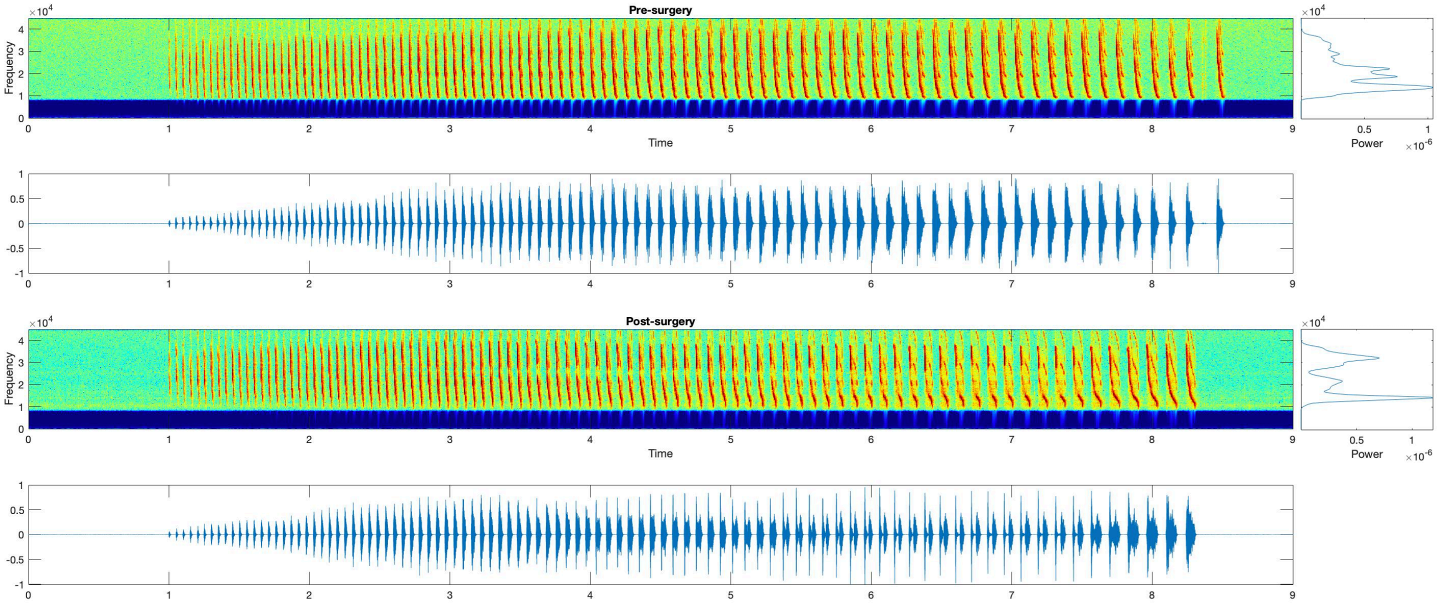


**Figure S7.** Spectrograms and oscillograms of a pre- (top) and post-surgery (bottom) song for an animal that had a minor ablation (individual E). The beginning of the post-surgery song is typical, but at approximately 3.5 seconds, frequencies in the middle of the range are lost.

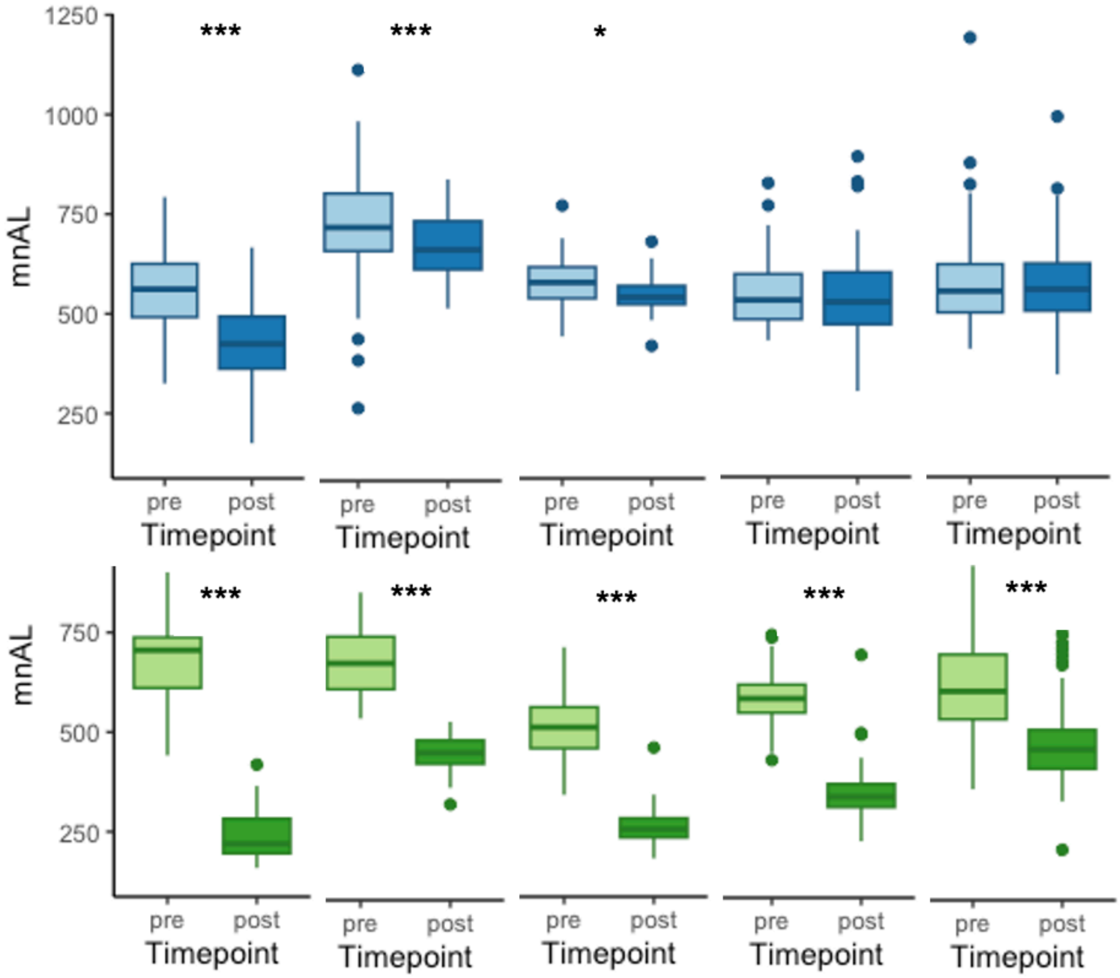


**Figure S8.** Boxplot of mean, normalized arc lengths for all songs of each individual in the pre- and post-surgery periods. Blue = sham, green = ablation. * $p < 0.05$, *** $p < 0.001$.

| | mean_nAL | | |
|---|---|---|---|
| *Predictors* | *Estimates* | *CI* | *p* |
| (Intercept) | 600.43 | 544.45 – 656.41 | **<0.001** |
| Treatment [ablation] | 1.23 | -82.94 – 85.39 | 0.977 |
| Timepoint [post] | -50.41 | -92.19 – -8.64 | **0.018** |
| Treatment [ablation] * Timepoint [post] | -164.77 | -228.28 – -101.26 | **<0.001** |

**Table S9.** Model results for dataset with individual A removed.
